# Defining functional neuraminidase inhibitor drug resistance motifs in avian influenza viruses and the consequential impact on virus fitness in chicken cells

**DOI:** 10.1101/663872

**Authors:** Dagmara Bialy, Holly Shelton

**Author notes:** Address correspondence to: Holly Shelton.

## Abstract

Neuraminidase inhibitors (NAIs) are antiviral agents recommended worldwide to treat or prevent influenza virus infections in humans. Mapping of functional resistance to currently licensed NAIs has been limited to human influenza viruses with only sporadic reports investigating avian influenza viruses (AIV). However past pandemics as well as the increasing number of humans infected with AIV have shown the importance of having information about avian NAs that could cross the species barrier. In this study we introduced four NAI resistance-associated mutations previously found in human strains into the NA of six prevalent AIV subtypes that threaten the poultry industry and human health: H7N9, H6N1, H4N6 and highly pathogenic H5N8, H5N6 and H5N2. Using the established MUNANA assay we show that R292K substitution significantly impaired NA activity in all strains, whereas E119V, H274Y and N294S have more variable effects on NA activity. The impact of these mutations on NAI susceptibility was drug- and strain-specific. We have shown that despite compromised NA activity drug-resistant H5N6 and H6N1 viruses replicated to comparable or significantly higher titres in primary chicken cells as compared to wild type. The replicative fitness of NAI-resistant H5N6 was also confirmed *in ovo*. Two drug resistant H5N6 viruses had single amino acid substitutions in their haemagglutinin (HA) which reduced receptor binding properties. Our results demonstrate that there are no universal NAI resistance determinants for all strains and although some are clearly deleterious for the virus, others can be rapidly compensated by acquiring concurrent changes in other gene segments.

**IMPORTANCE:** The number of human infections caused by avian influenza viruses (AIV) keeps increasing. This together with the rapid emergence of influenza strains resistant to neuraminidase inhibitor drugs (NAIs) observed in the past raises a significant concern to public health. We studied the NAI resistance-associated molecular changes, previously reported in neuraminidase (NA) of human influenza, in AIV background. We found that single amino acid substitution can confer a multidrug resistance, or lead to a single-drug resistance across multiple virus subtypes. We also found that the drug-resistant viruses retained or showed enhanced fitness properties as compared to the corresponding wild-type, and this could be achieved by quick acquisition of concurrent mutations in haemagglutinin. Our study highlights the need for constant monitoring of NAI-resistance in AIV and understanding the molecular basis of antiviral resistance, as such information would be invaluable for pandemic preparedness and may facilitate the development of novel therapeutics.

## INTRODUCTION

Wild aquatic birds are the major reservoir of avian influenza virus (AIV) worldwide. AIVs frequently transmit from these reservoir species to other bird populations such as domestic poultry, from where they can cross additional species barriers and jump into mammalian hosts such as humans. An increase of human infections caused by different AIV subtypes in the recent years [1–6] together with the evidence pointing at the avian origins of past influenza pandemics [7, 8] places AIV strains as a significant concern to public health.

The emergence of drug resistance is an inherent risk associated with therapeutic use of antivirals. There are three virus targeted classes of drugs available for treatment of influenza virus infections in humans: 1) adamantanes (amantadine and rimantadine) – blocking the function of the M2 ion channel protein in influenza inhibiting uncoating of the virus particle; 2) viral polymerase inhibitors (pimodivir, favipiravir and baloxavir acid) – modulation of viral replication, currently approved only in certain countries or in late stage clinical trials; and 3) neuraminidase inhibitors (NAIs) - targeting the sialidase active site of influenza neuraminidase (NA) which acts to limit virus spread and transmission. Adamantanes are no longer recommended for use in humans since drug-resistant seasonal H3N2 and H1N1 currently circulate and resistance in AIV strains such as H5N1 has also been widely reported [9–11].

With significantly limited effectiveness of adamantanes, NAIs are currently recommended by WHO for the first-line treatment and prophylaxis of influenza infections in humans. There are four currently licensed NAIs: oseltamivir (OSE), zanamivir (ZAN), peramivir (PER) and laninamivir, these drugs differ in their chemistry and administration route, but all similarly to amantadine can support the selection of drug-resistant viruses if used in an uncontrolled manner or for prolonged periods of time [12–15]. During the 2007-2008 influenza season the majority of circulating human H1N1 isolates were found to be resistant to OSE suggesting that the resistant viruses which were antigenically similar to OSE-sensitive variants had a selective advantage [16, 17]. However importantly, the advantage prescribed by the resistant viruses was not driven by selective drug pressure rather other concurrent fitness motifs which were associated with the drug resistant viruses [18, 19]. The functional resistance to NAI drugs is not commonly observed in circulating AIVs however it emerges rapidly in human patients infected with AIV and treated with antivirals as it was documented in cases of H7N9 or H5N1-infected individuals [14, 15].

The molecular signatures of NAI resistance have been broadly studied in human N1 and N2 and are linked but not limited to four most commonly observed changes in the NA: three occurring at the framework residues (E119V, H274Y, N294S) and one alteration within the catalytic site (R292K) [12, 16, 17, 20, 21]. Despite the increasing number of human infections caused by AIVs there are very few reports investigating the molecular basis or the fitness costs incurred by acquisition of NAI resistance in avian strains. This knowledge gap is significant considering the chance that there is the potential for the next influenza pandemic virus to carry an avian NA genetic segment either as the result of wholly avian origin crossing the species barrier or via reassortment.

In this study we introduced four NAI resistance-associated mutations (E119V, H274Y, R292K, N294S) into five different avian NA subtypes, N1, N2, N6, N8 and N9 in the context of current AIVs which have caused infections in humans (H5N6, H7N9 and H6N1) or have evident zoonotic potential (H5N8, H5N2 and H4N6). We determined the susceptibility of recombinant viruses to three licensed NAIs: OSE, ZAN and PER and the impact of introduced changes on viral NA activity. Furthermore, we assessed virus fitness of selected drug-resistant strains in primary chicken kidney cells (cKC) as well as in embryonated hen’s eggs, and dissected the molecular mechanism accounting for their growth kinetics as compared to viruses carrying a wild type (wt) NA.

## RESULTS

### Prevalence of human influenza virus resistance associated mutations in avian influenza virus strains

To investigate the molecular signatures of NAI-resistance in various AIV NA backgrounds we chose a panel of prevalent avian influenza strains posing a current threat not only to poultry and the associated industries, but also to human health due to their zoonotic potential (Table 1).

**Table 1.** Panel of generated recombinant influenza strains (2:6) HxNx : PR8. * – human AIV isolates

| Group | Subtype | HA/NA | Strain | Hazard to poultry and public health | Prevalence of NAI resistance-associated mutations in AIVs 2013-2018 (E119V/D, H274Y, R292K, N294S) |
| --- | --- | --- | --- | --- | --- |
| N2-like | **N2** | **H5N2**<br>HPAIV<br>clade<br>2.3.4.4 | A/goose/Taiwan/01031/2015 | Serious concern to poultry industry, with the 2014/15 outbreak in North America only affecting >50 million birds [24]. | none/438 |
|  | **N6** | **H5N6**<br>HPAIV<br>clade<br>2.3.4.4 | A/chicken/Jiangxi/02.05<br>YGYXG023-P/2015 | 23 confirmed human cases; 15 deaths (as of November 2018 since 2014) [4, 5]. | H274Y – 1/1041 (0.1%)<br>N294S – 3/1041 (0.3%)<br>E119D – 10/1041 (1%)<br>*E119D - 2/22 (10%) |
|  |  | **H4N6** | A/chicken/Hunan/S1267/2010 | Preferential binding of swine H4N6 isolates to human receptors; further evidence for ongoing mammalian adaptation [30]. | none/230 |
|  | **N9** | **H7N9** | A/Anhui/1/2013 | 1568 confirmed human cases; 615 deaths (as of October 2018 since 2013) [6]. | none/743<br>*E119V – 5/1233 (0.4%)<br>*H274Y – 4/1233 (0.3%)<br>*R292K – 45/1233 (3.6%)<br>*N294S – 1/1233 (0.08%) |
| N1-like | **N8** | **H5N8**<br>HPAIV<br>clade<br>2.3.4.4 | A/scarlet ibis/Germany/Ar44-L01279/2015 | Multiple outbreaks with lethal outcomes in avian species in Africa, America, Asia, Europe and Middle East since 2014 [25]. | none/623 |
|  | **N1** | **H6N1** | A/chicken/Taiwan/67/2013 | First human infection confirmed in Taiwan in 2013 [1]. | none/40 |

\* – human AIV isolates

We selected three contemporary highly pathogenic avian influenza viruses (HPAIV) representing clade 2.3.4.4: H5N6, H5N8 and H5N2, all rapidly spreading across the globe and causing multiple die-offs in wild migratory birds as well as significant outbreaks in domestic poultry. To date human infections have been confirmed only for H5N6 subtype, with a total of 23 cases and at least 15 deaths reported in China since 2014 [4, 5, 22–25].

In addition we included three low pathogenic avian influenza strains including H7N9 which emerged in China in 2013 infecting poultry species widely in that geography and spilling over into humans. The total number of H7N9 human cases has exceeded 1500 with a case fatality rate of ∼39% [2, 6]. H6N1 became a concern to public health in June 2013 when the first human infection was reported in Taiwan [1]. This subtype represents a great zoonotic potential due to the ongoing genetic changes altering receptor binding specificity [26, 27], and remains highly prevalent in Taiwanese poultry species [28]. Recent serological evidence implies that agricultural workers exposed to diseased poultry during an unrecognized outbreak may have been asymptomatically infected with H4N6 avian influenza viruses [29]. Furthermore, genetic studies suggest that avian H4N6 strains are subject to frequent reassortment events, and swine H4N6 isolates show strong evidence of ongoing mammalian adaptation leading to a receptor binding switch towards human host [30, 31]

The four most common NAI resistance associated mutations in human influenza strains were E119V, H274Y, R292K and N294S. We assessed the prevalence of these four mutations in avian isolated and where available, human isolated AIV sequences in the subtypes we selected. We analysed the sequences that had been deposited in GISAID database between 2013, when the first human infections with avian H5N6, H7N9 and H6N1 were reported, and 2018. We identified H274Y substitution in 0.1% and N294S - in 0.3% of the total of 1041 avian H5N6 isolates (Table 1). None of the avian or human H5N6 carried E119V signature, however 10% of human and 1% of avian strains contained D at this position, a mutation recently reported to confer pan-resistance to available NAIs in pH1N1 [32].

### Generation of recombinant influenza strains bearing NAI-associated mutations

We introduced the described 4 point mutations as individual mutations into the NA genes of all six subtypes using site-directed mutagenesis and rescued the recombinant viruses with 6 internal genes of egg-adapted PR8 virus (PB2, PB1, PA, NP, M and NS) and matching HA/NA set of the particular AIV strain. We pre-selected the four NAI resistance-associated molecular signatures, previously identified in a number of human IAV isolates including H1N1, H3N2 and H7N9, and shown to confer resistance to at least one NAI drug [12, 13, 21, 33–38]. One substitution occurs at the actual catalytic residue (R292K) critical for the enzymatic function of NA, whereas three other changes (E119V, H274Y and N294S) are located at the framework positions stabilizing the structure of the active site of NA (Figure 1).

**Figure 1.**
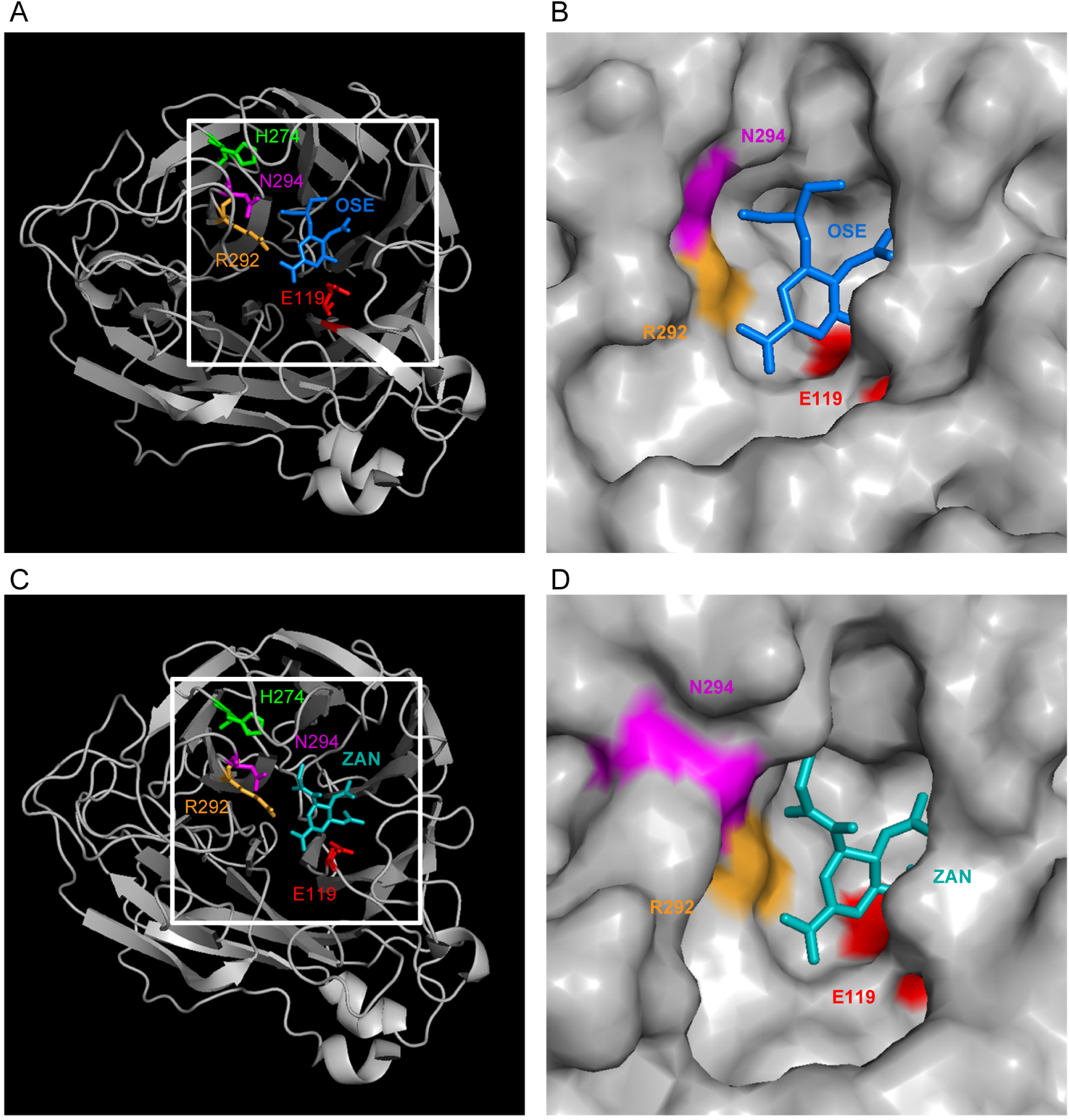
Location of potential neuraminidase inhibitor (NAI) resistance mutations in NA catalytic site. Four residues critical for binding of neuraminidase inhibitor drugs – three framework (E119, H274, N294) and one catalytic (R292) are highlighted in colours. Oseltamivir (OSE; A & B) and zanamivir (ZAN; C & D) drugs are shown in blue. **(A)** Crystal structure of H7N9 A/Anhui/1/2013 neuraminidase (NA) complexed with OSE was adapted from PDB 4MWQ. **(B)** Surface area of OSE binding pocket within the NA active site. **(C)** Crystal structure of H11N6 A/duck/England/1/1956 NA complexed with ZAN (PDB 2CML). (D) Surface area of ZAN binding cavity within the NA catalytic site.

All viruses were successfully rescued and passaged in embryonated hens eggs to generate working stocks, although the titres obtained for some of the mutant viruses were lower than the wild-type NA-carrying viruses (Supplementary table 1). The introduced NA mutations were maintained upon rescue and passage with no additional mutations in the NA gene appearing.

### Impact of introduced mutations on neuraminidase activity

Following introduction of NAI resistance-associated mutations into viral NA segment we assessed the sialidase activity of recombinant viruses using MUNANA assay. The NA activities were expressed as the amount of released fluorescent product, 4-MU (4-methylumbelliferone), per 1 ml of virus suspension within 1 hour at 37°C. These values were normalised to haemagglutination titre (HAU) / of the viral stock and displayed as a fold difference compared to the reference strain, PR8 (Figure 2). We found that all viruses bearing wt NA except H4N6 showed an increased NA activity/HAU ratio compared to PR8. The R292K substitution, located within the catalytic site of the enzyme, significantly compromised NA activity of all recombinant viruses when compared to the corresponding wt (Figure 2A-F; orange bars). The NA activity/HAU ratio of H5N8_R292K mutant could not be determined as the virus exhibited no detectable haemagglutinating properties on chicken red blood cells. This virus was however able to generate plaques on an MDCK cell layer.

**Figure 2.**
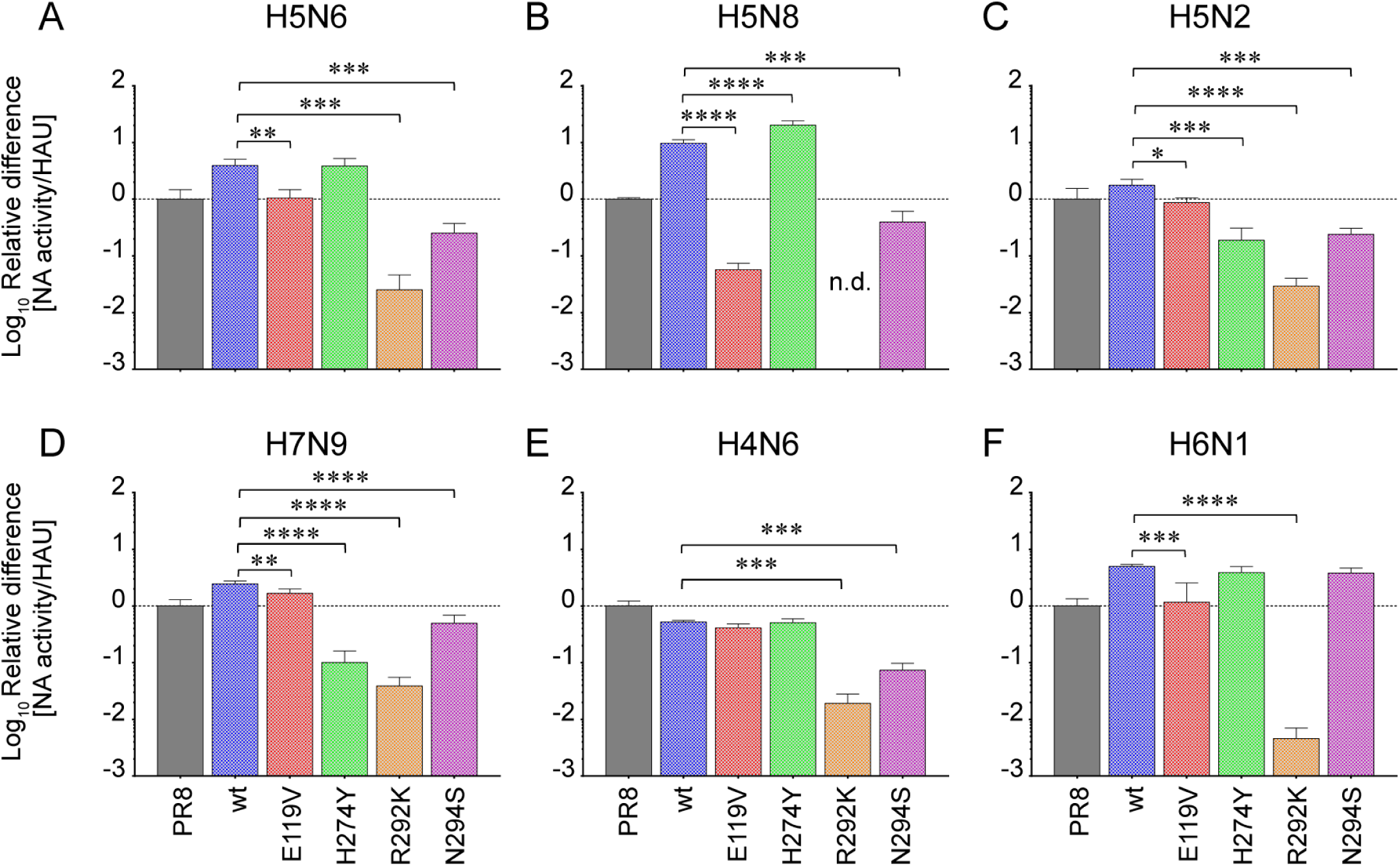
Impact of introduced mutations on NA activity of recombinant strains. Neuraminidase (NA) activities of six virus subtypes: H5N6 (A), H5N8 (B), H5N2 (C), H7N9 (D), H4N6 (E) and H6N1 (F) were determined using MUNANA assay. Equivalent NA activity levels (corresponding to known amount of 4-MU/ml released per hour at 37°C) were chosen for a panel of 5 viruses representing each subtype (wt, E119V, H274Y, R292K, N294S). These values were then normalised to HAU of virus stock, and the ratio NA activity/HAU for a reference strain (PR8) was set to 1 and is indicated by a horizontal dashed line on each graph. The Y axis shows log_10_ fold difference of NA activity/HAU between PR8 and tested viruses. Data represent mean of 3 independent experiments ± SD; statistical significance was determined by one-way ANOVA and Dunnett’s multiple comparisons tests (* P<0.05; ** P<0.01; *** P<0.001).

The impact of three remaining mutations on the enzymatic activity of NA varied among virus subtypes. The E119V mutation (red bars) significantly reduced the NA activity of H5N6, H5N8, H5N2, H7N9 and H6N1 (Figure 2A, B, C, D & F, respectively), but not of H4N6 (Figure 2E). The substitution H274Y (green bars) led to a significant drop in NA activity for H7N9 and H5N2, and an increase for H5N8, whereas other subtypes remained unaffected. The N294S amino acid change (magenta bars) significantly impaired NA activity of all subtypes except H6N1.

To determine the relative NA activity of H5N8_R292K we normalised the raw values to PFU (Plaque Forming Units) and compared the ratios for H5N8 subtype (Supplementary Figure 1B). The NA activity/PFU of H5N8_R292K mutant was significantly reduced (∼6000 times) compared to wt NA. E119V and N294S led to a significant reduction whereas H274Y substitution resulted in a significant increase of sialidase activity of recombinant viruses (Supplementary Figure 1). These results corresponded to the NA activity profile with values normalised to HAU. (Figure 2B)..

### Susceptibility profile of tested viruses to licenced neuraminidase inhibitors (NAIs)

We next determined the susceptibility of the generated viruses to three neuraminidase inhibitor drugs: oseltamivir (OSE), zanamivir (ZAN) and peramivir (PER) using the MUNANA fluorescence-based NA inhibition assay to generate IC_50_ values. IC_50_ is the concentration of drug that will reduce the NA activity by 50% from no drug (Figure 3 and Table 2). The outcomes of the assay were interpreted using the WHO recommendations for classification of antiviral drug resistance: normal inhibition (NI; <10-fold increase in IC_50_ compared to a corresponding drug-sensitive strain), reduced inhibition (RI; IC_50_ increase between 10 – 100-fold), highly reduced inhibition (HRI; >100-fold increase).

**Figure 3.**
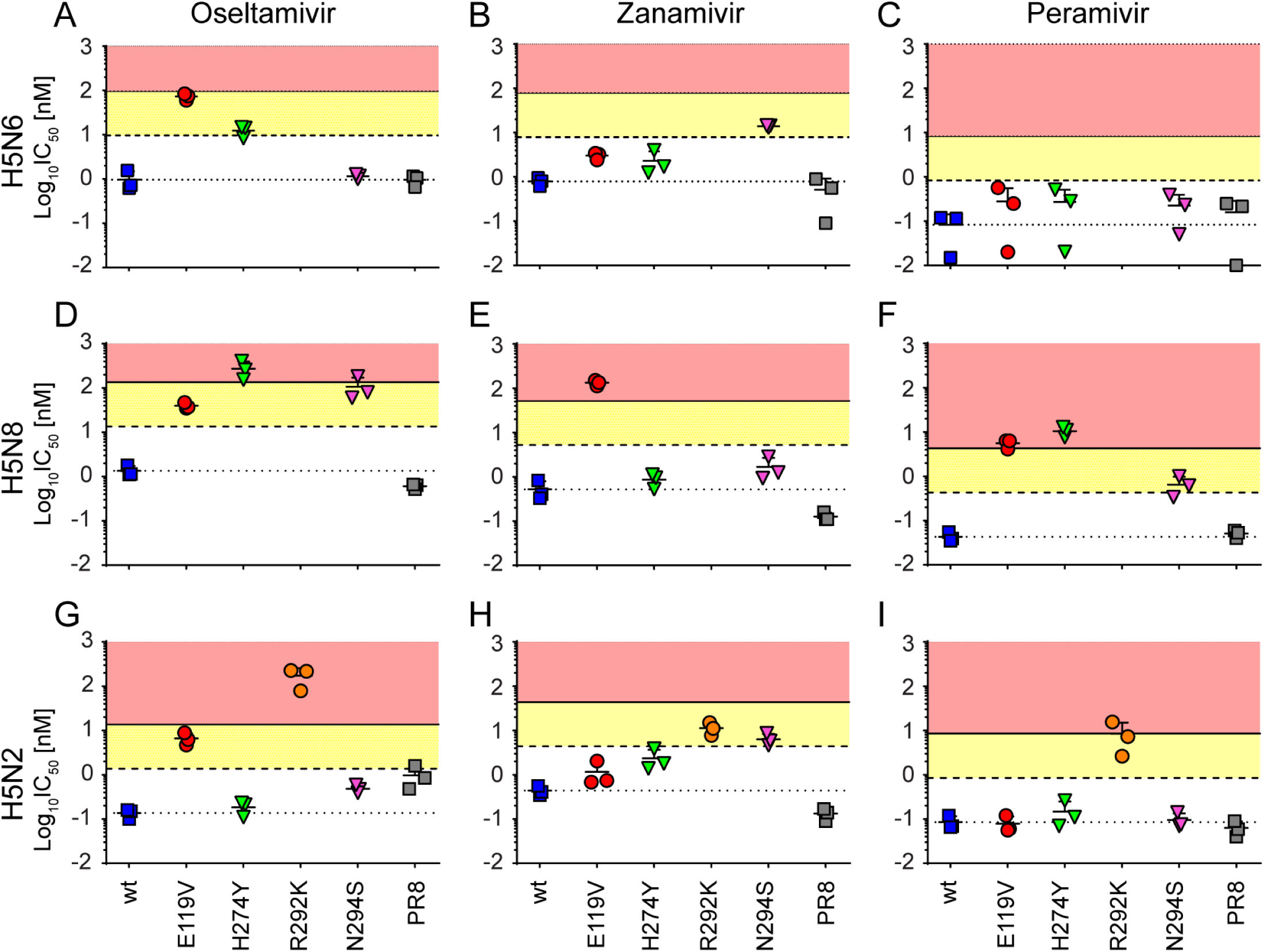
Neuraminidase inhibitor (NAI) susceptibility profile of H5N6, H5N8 and H5N2 influenza viruses as determined by MUNANA assay. The IC_50_ values for three HPAI (highly pathogenic avian influenza): H5N6 (A – C), H5N8 (D – F) and H5N2 (G – I) were determined by fluorescent NA inhibition assay. Each panel of viruses (R292K only for H5N2) alongside with the PR8 strain were tested for their susceptibility to OSE (left column), ZAN (middle column) and PER (right column). Short horizontal line with error bar indicates the mean of log_10_IC_50_ [nM] ± SD of three independent experiments; each data point represents a single experiment. The area between the dotted and dashed lines on each graph represents the normal inhibition (NI; <10-fold increase in IC_50_ over corresponding wt), the yellow-shaded area - reduced inhibition (RI; 10 - 100-fold increase in IC_50_) and the red-shaded area – highly reduced inhibition (HRI; 100<-fold increase in IC_50_).

**Table 2.**
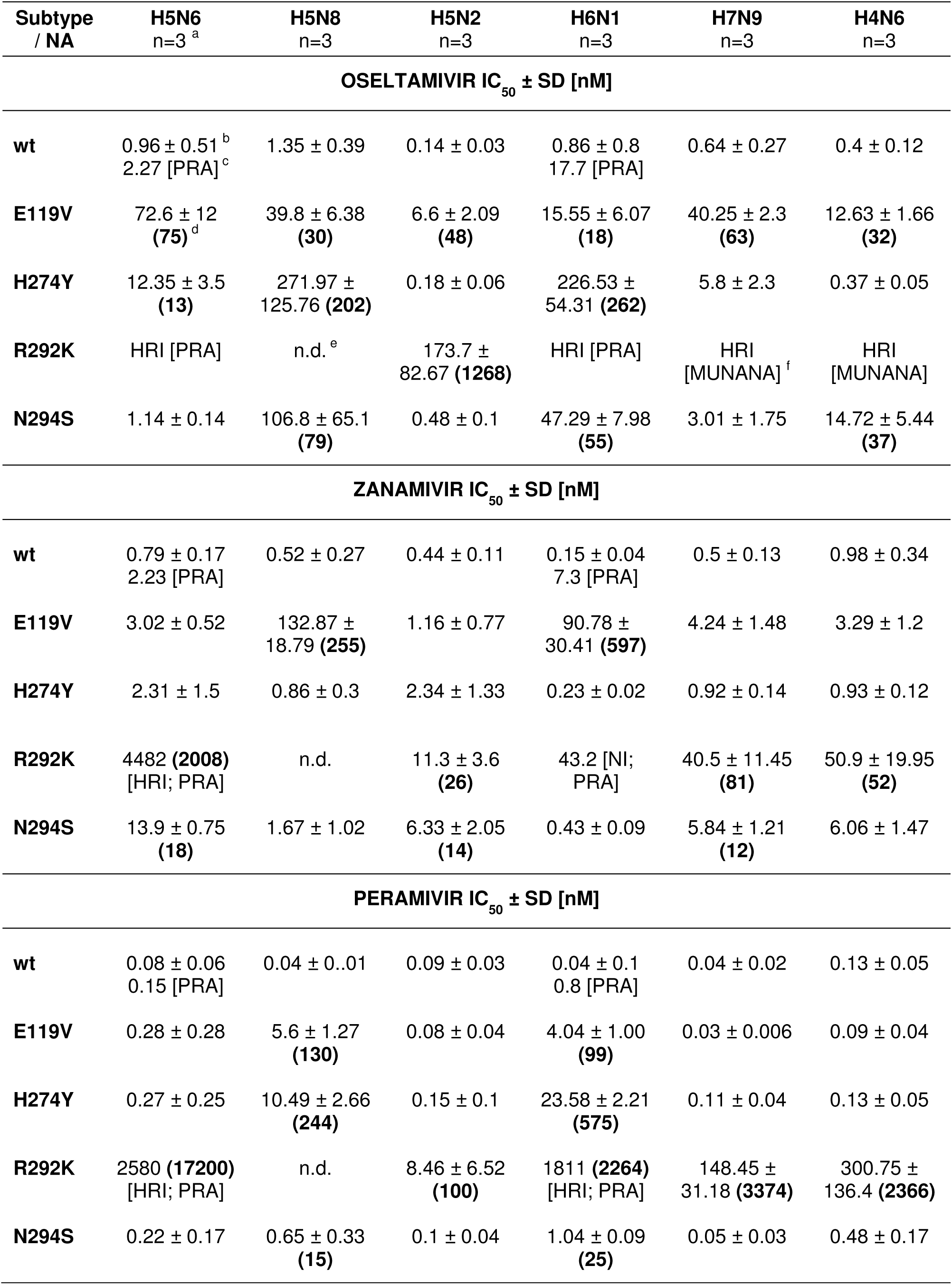
The impact of introduced mutations on virus susceptibility to licensed NAI drugs in different HA/NA backgrounds. a – number of independent MUNANA assays b – data represent mean of n independent experiments ± standard deviation (SD) c – drug susceptibility determined by <u>P</u>laque <u>R</u>eduction <u>A</u>ssay d – numbers in parentheses indicate fold increase in IC_50_ over the corresponding wt NA; shown only for reduced inhibition (RI; 10 - 100-fold) or highly reduced inhibition (HRI; >100-fold) e – not determined f – drug susceptibility determined by MUNANA assay, although dose response curve not converged (no precise IC_50_ could be retrieved)

The graphic representation of NAIs IC_50_ shows that the E119V and H274Y mutations introduced in the NA of H5N6 background resulted in RI by OSE (75- and 13-fold increase, respectively; Figure 3A), and that N294S led to RI by ZAN (18-fold increase; Figure 3B). All three substituted NA variants remained susceptible to PER (Figure 3C). The IC_50_ values for H5N6_R292K were determined using plaque reduction assay (PRA), as the fluorescent signal generated using MUNANA did not reach the recommended threshold for signal-to-background ratio. No reduction in plaque size was observed for H5N6_R292K even in the presence of the highest OSE concentration (10 μM; Figure 4A and D), whereas the same concentration of ZAN or PER reduced the H5N6_R292K plaque size by approximately 40% compared to wt (Figure 4B and E; C and F, respectively). Based on the generated dose response curves of the PRA we concluded that H5N6_R292K showed HRI for all three NAIs: it remained unresponsive to OSE treatment and showed a 2008-fold IC_50_ increase over wt for ZAN and 17200-fold IC_50_ increase for PER (Table 2 and Figure 4).

**Figure 4.**
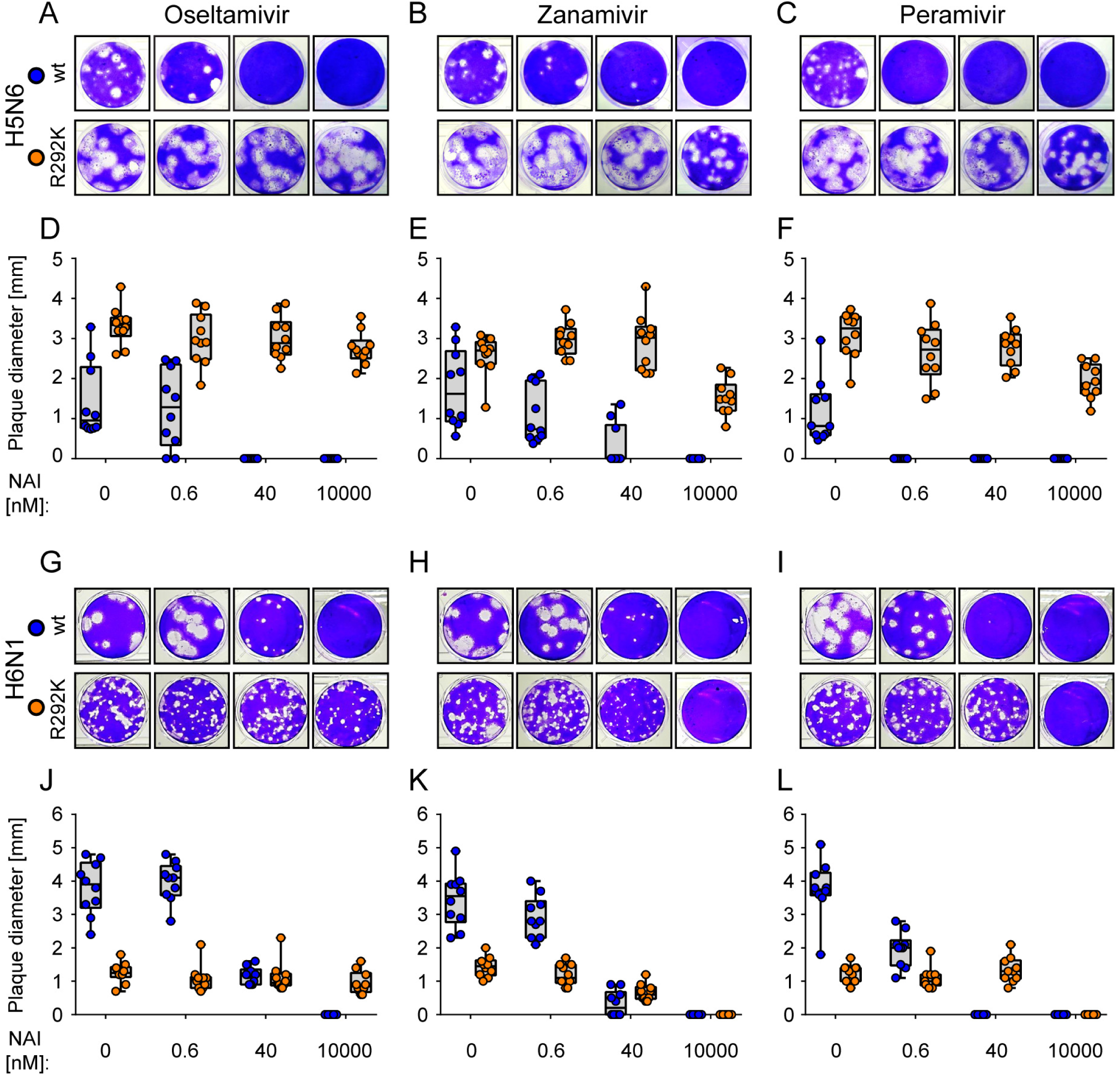
Susceptibility profiles to NAIs of H5N6 and H6N1 viruses carrying R292K neuraminidase substitution as determined by plaque reduction assay. MDCK cells were infected with H5N6 (A-F) or H6N1 (G-L) viruses bearing either wt or R292K neuraminidase variants. The cells were cultured at 37°C with 0.6% Avicel overlay and increasing NAI concentrations (0 – 10 μM): OSE (left column), ZAN (middle column) or PER (right column). The plaques were developed after 72 h of incubation (A-C; G-I) and the plaque sizes were measured for 10 plaques per condition (D-F; J-L). Box and whiskers graphs in D-F & J-L represent the median plaque diameter with minimum and maximum values [mm]; each data point represents a single plaque.

The E119V substitution tested in the H5N8 background conferred resistance to all three inhibitors, with HRI for ZAN and PER (255-fold and 130-fold increase, respectively; Figure 3E - F) and RI for OSE (30-fold increase; Figure 3D). The viruses bearing H274Y and N294S showed IC_50_ increase for both, OSE (202-fold and 79-fold, respectively; Figure 3D) and PER (244-fold and 15-fold; Figure 3F), but not ZAN (Figure 3E). The drug resistance profile for H5N8_R292K could neither be determined by MUNANA assay because of insufficient NA activity, nor PRA, due to small size of the plaques formed by wt H5N8, making visualisation of the control virus unreliable.

The R292K catalytic site substitution in the H5N2 backbone led to HRI by OSE (1268-fold increase; Figure 3G) and RI by ZAN and PER (26- and 100-fold increase, respectively; Figure 3H - I). Two other mutations induced an increase in IC_50_ for a single drug each: E119V for OSE (48-fold; Figure 3G) and N294S for ZAN (14-fold; Figure 3H). The H5N2_H274Y remained sensitive to all three antivirals tested (Figure 3G - I).

As determined by MUNANA assay three of the mutations introduced in H6N1 background caused resistance to OSE (E119V 18-fold, H274Y 262-fold and N294S 55-fold increase) and PER (E119V 99-fold, H274Y 575-fold, and N294S 25-fold increase), whereas only one substitution reduced the susceptibility to ZAN (E119V, 597-fold increase in IC_50_). Drug resistance profile for H6N1_R292K was determined using PRA due to very low NA activity in MUNANA assay and gave an IC_50_ value 2264-fold higher for PER (Figure 4I & L). Although the IC_50_ for OSE could not be retrieved from the dose-response curve of the PRA (no reduction in plaque size was observed in the presence of drug), the virus was classified as highly resistant to this inhibitor since no inhibition of replication was observed even at the highest concentration of drug tested (10000 nM) (Figure 4G & J).

One amino acid change in H7N9 virus induced high resistance to PER (R292K, 3374-fold increase), whereas two substitutions – R292K and N294S - conferred RI by ZAN (81- and 12-fold increase, respectively). H7N9_E119V was shown to be resistant to OSE (63-fold increase). Although the IC_50_ for H7N9_R292K could not be determined from OSE dose-response curve of MUNANA assay, the virus was classified as highly resistant to this drug as no reduction in fluorescent signal was detected even at the highest (10 μM) OSE concentration.

Reduced inhibition to OSE was found when E119V and N294S substitutions were placed in H4N6 background (32- and 37-fold increase in IC_50_, respectively). H4N6_R292K virus exhibited 52-fold increase in IC_50_ for ZAN (RI), 2366-fold increase for PER (HRI), and high resistance level to OSE as no reduction in fluorescence signal was observed in the presence of increasing drug concentrations.

In summary, we screened 30 avian viruses and identified 15 viruses resistant to OSE (of which 5 showed HRI), 9 to ZAN (3 were HRI) and 11 to PER (7 were HRI) (Table 2).

### In vitro replication kinetics of NAI-resistant H5N6 and H6N1 viruses in cKC

To understand whether introduced mutations, besides compromising NA activity, also affect virus ability to support efficient infection, we used primary chicken kidney cells (cKC) and assessed the replicative capacity of two AIV subtypes bearing NAI resistance-associated markers. The viruses were selected based on the following criteria: 1) the mutations introduced in NA gene conferred functional resistance to at least one NAI, as determined in NAI inhibition assay (Table 2); 2) the mutant viruses grew to comparable titres as the corresponding wt parental viruses *in ovo*; (Supplementary table 1); 3) the mutant viruses were able to spread in cell monolayer and form plaques of well-defined morphology, as determined during plaque assay titrations (Supplementary figure 3). We selected wt H5N6 alongside three of the isogenic NAI-resistant mutants: E119V - highly resistant to OSE, R292K – highly resistant to all three NAIs tested in this study, and N294S - resistant to ZAN only. The H5N6_H274Y, although resistant to ZAN, did not yield comparable titres after propagation in eggs to the wt virus. The second set of viruses included wt H6N1 and three NAI resistant variants: E119V – resistant to all three NAIs with high resistance to ZAN, H274Y and N294S – both resistant to OSE and PER, with H274Y showing high resistance to both drugs. H6N1_R292K, although highly resistant to OSE and PER, was not included in the assay due to low virus titres produced in eggs and attenuated virus spread *in vitro*, as indicated by small plaque phenotype in MDCK cells (Figure 4 G-I).

We inoculated primary cKC at an MOI of 0.001 PFU/cell with each virus in triplicate and collected the supernatants at the indicated time points (Figure 5A-B). All three NAI-resistant H5N6 viruses as well as the wt showed the peak of viral replication at 48 h p.i. and the titre dropped by ∼0.5-2 logs at 72 h p.i. (Figure 5A). When compared to wt H5N6, all three mutant viruses grew to significantly higher titres at 8 h (E119V p < 0.01; R292K p < 0.01; N294S p < 0.001), 24 h (E119V p < 0.0001; R292K p < 0.0001; N294S p < 0.0001), 48 h (E119V p < 0.0001; R292K p < 0.0001; N294S p < 0.0001) and 72 h post infection (E119V p < 0.0001; R292K p < 0.0001; N294S p < 0.0001). Addition of OSE to cKC infected with wt H5N6 significantly reduced the titre of cell-free virus at 24, 48 and 72 h p.i. when compared to untreated cells (all p < 0.0001). No infectious virus particles were detected in OSE-treated samples at 8 h p.i. by plaque assay (p < 0.05). Cell viability remained unchanged upon addition of drug to cKC cultures (Supplementary Figure 2).

**Figure 5.**
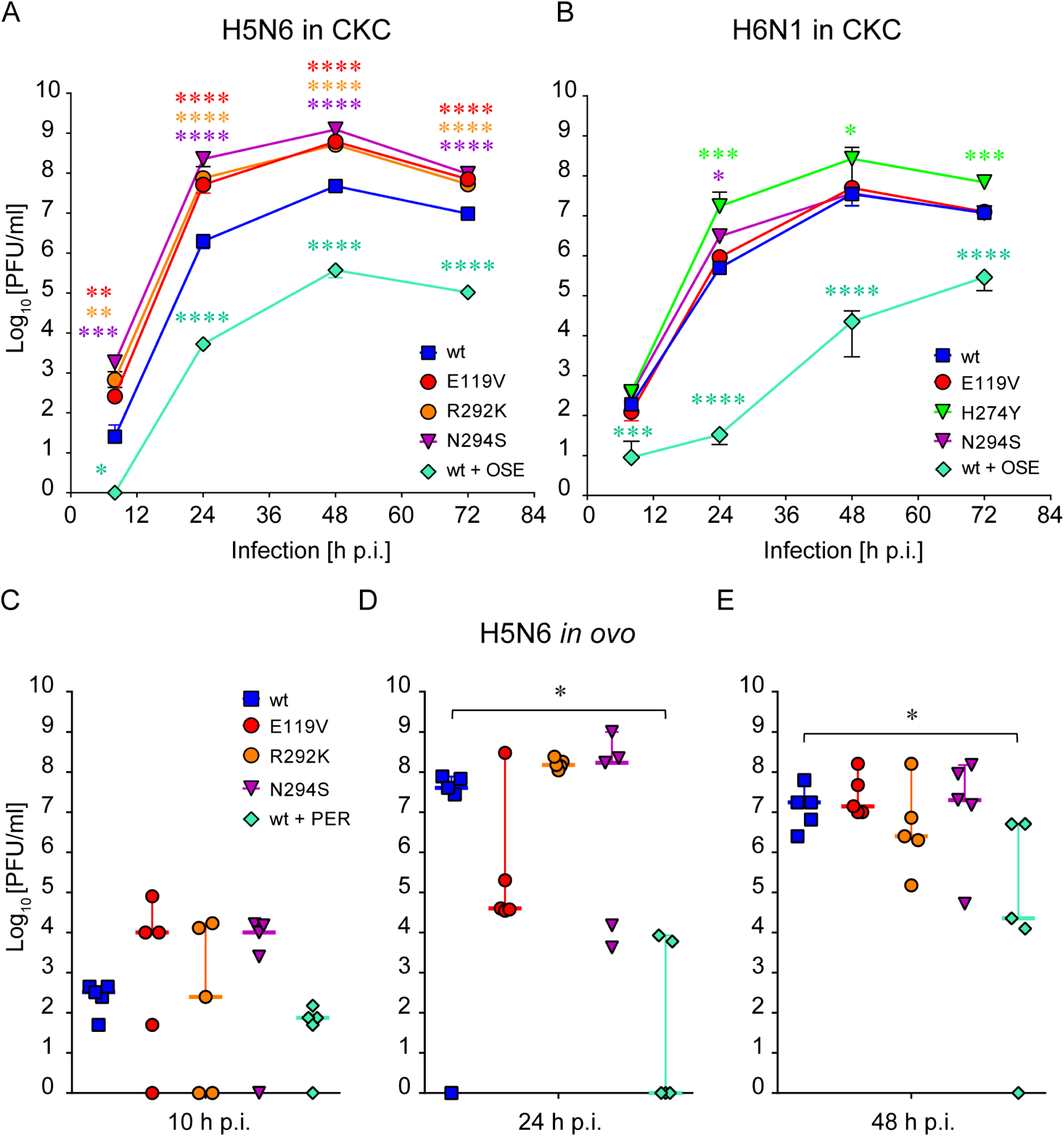
Growth kinetics of NAI-resistant H5N6 and H6N1 viruses *in vitro* and *in ovo*. **(A - B)** cKC were infected with H5N6 (A) or H6N1 (B) NAI-resistant viruses as well as the corresponding wt at an MOI of 0.001. The supernatants were collected at the indicated time points and titrated by plaque assay. Graphs shown are representative of 2 independent experiments; each data point represents mean ± SD; n = 3. Statistical significance was analysed using one-way ANOVA and Dunnett’s multiple comparisons tests (* P<0.05; ** P<0.01; *** P<0.001). **(C - E)** 10 day-old SPF embryonated hen’s eggs were inoculated with H5N6 viruses at 100 PFU/egg. Allantoic fluid samples were collected at the indicated time points and titrated. Each data point represents a single egg; short horizontal line indicates the median with 95% CI, n = 5. All virus-infected embryos were viable at 10 and 24 h p.i. and reached the terminal end point by 48 h p.i. Statistical significance was determined by one-way ANOVA and Dunnett’s multiple comparisons tests (* P<0.05; ** P<0.01; *** P<0.001).

Similar to H5N6, all H6N1 viruses reached the peak of viral replication at 48 h p.i., and their titres dropped at 72 h p.i (Figure 5B). H6N1_H274Y replicated to significantly higher titres at 24 h (p < 0.001), 48 h (p < 0.05) and 72 h p.i. (p < 0.001) as compared to wt. H6N1_E119V and N294S viruses grew to comparable titres as the wt throughout the time course, with N294S producing slightly higher titre at 24 h p.i. (Figure 5B; p < 0.05). Similarly to H5N6, addition of OSE to cKC cultures infected with wt H6N1 significantly reduced viral titres throughout the time course (8 h p.i. - p < 0.001; 24, 48 and 72 h p.i. – p < 0.0001).

Taken together the results of virus replication kinetics in cKC have shown that the NAI-resistant viruses, despite compromised sialidase activity, replicated to similar or significantly higher titres when compared to the corresponding drug-sensitive viruses bearing wild-type NAs.

To confirm the effects we observed in cKC we inoculated 10-day old embryonated hen’s eggs with drug-resistant H5N6 viruses, collected the samples of allantoic fluid at indicated time points and determined the median titres of infectious virus (Figure 5 C – E; five eggs per virus and per time point). At 10 and 24 h p.i. all virus-infected embryos remained alive. At 10 h p.i. the median titres of H5N6_E119V and N294S viruses reached ∼1 × 10^4^ PFU/ml, which was ∼2-log more than wt H5N6 and R292K (Figure 5C). At 24 h p.i. wt H5N6 together with two NAI-resistant mutants: R292K and N294S, replicated to their highest titres between 4 × 10^7^ – 1.5 × 10^8^ PFU/ml (Figure 5D), and the numbers of infectious particles dropped at 48 h p.i. (Figure 5E), when all the embryos reached their terminal end points as indicated by extensive haemolysis of supporting blood vessels. In contrast to other viruses, E119V showed a mild increase in titre between 10 and 24 h p.i. (from 1 to 4 × 10^4^ PFU/ml) and reached the maximum of 1.4 × 10^7^ PFU/ml only at 48 h p.i.

Pre-incubation of wt H5N6 with PER reduced viral titres when compared to eggs inoculated with wt H5N6 only (10 h - 0.5-log reduction; 24 h - 7-log reduction, p < 0.05; and 48 h p.i. - 3-log reduction, p < 0.05), but it was not sufficient to completely abrogate virus production. The differences in virus growth in eggs observed for wt H5N6 and the corresponding NAI-resistant variants were not significant, thus based on the results of our *in vitro* and *in ovo* fitness assessment we concluded that the NAI-resistant H5N6 viruses, despite reduced NA activity, are able to replicate at least as efficiently as the wild-type in tested models.

### Compensatory mutations in HA of NAI-resistant H5N6 and their impact on receptor binding avidity

Concurrent mutations in HA can re-balance changes affecting NA activity therefore we analysed the HA sequences of NAI-resistant H5N6 and H6N1 viruses. While no changes in HA of H6N1 were found, two H5N6 mutants acquired single amino acid substitutions in their HA following one passage in eggs: Y98F (H3 numbering) in the NA R292K mutant and A189T in N294S NA mutant virus (Figure 6A). Both mutations have previously been shown to alter receptor binding affinity in human strains [19, 39]. We used partly desialylated cRBCs to explore the impact on receptor binding avidity in H5N6 of the Y98F and A189T mutations in HA (Figure 6B). Both HA mutants showed significantly reduced receptor binding avidity as compared to wt: the Y98F HA mutation present in the R292K NA virus was reduced by 94% and the A189T HA mutation present in the N294S was reduced by 95% (both p < 0.01). Surprisingly, the NA E119V mutant, despite lack of compensatory mutations in HA also exhibited decreased receptor binding avidity by ∼55% (p < 0.05).

**Figure 6.**
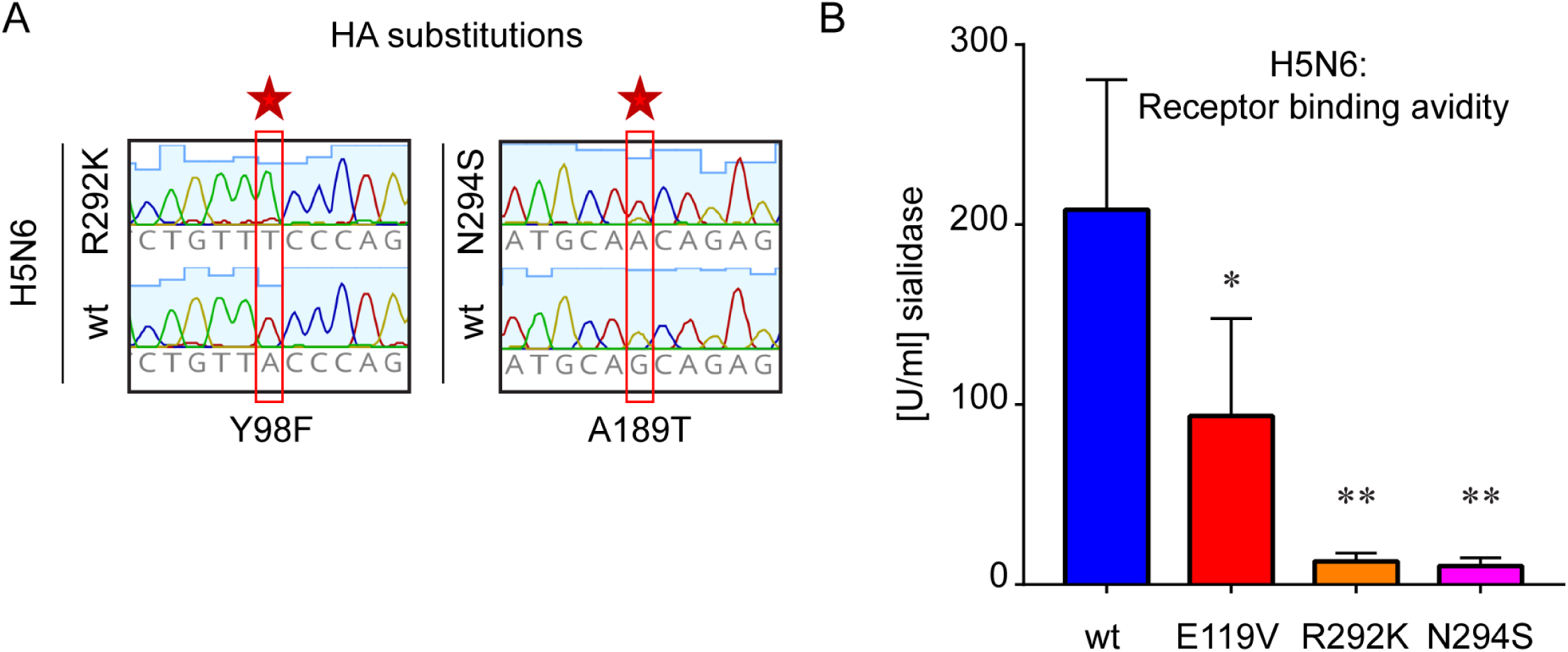
Concurrent changes in HA of NAI-resistant H5N6 viruses and their impact on receptor binding avidity. (A) Amino acid substitutions found in HA genes of H5N6_R292K and N294S viruses. HA segments or their fragments were amplified from viral RNA and analysed following Sanger sequencing. Changes on nucleotide level are indicated with a red frame and a star; amino acid substitutions are shown below (H3 numbering). (B) Receptor binding avidity of H5N6 viruses. Relative receptor binding avidity was determined with 4 HAU of each virus on partially de-sialylated cRBCs pre-treated with recombinant NA from *C. perfringens*. The Y-axis shows mean ± SD of maximal NA concentration allowing for full haemagglutination of each virus (U/ml); n = 3. Statistical significance was analysed using one-way ANOVA and Dunnett’s multiple comparisons tests (* P<0.05; ** P<0.01).

## DISCUSSION

In our study we characterised the resistance profile of AIV strains to currently licensed NAIs and further assessed the fitness of selected drug-resistant variants. We introduced four NAI resistance-associated mutations in the NA gene of recombinant AIVs, all four mutations had previously been reported to confer resistance to licensed NAIs in human H1N1 and H3N2 strains [12, 16, 17, 20, 21, 38]. Analysis of the GISAID database revealed that only the H5N6 subtype of AIVs we investigated possessed any of the four mutations in sequences isolated from avian hosts and these were present at very low percentages (<0.5%; Table 1). Resistance signatures in human isolates are likely to arise in hospitalised patients with confirmed or suspected influenza, following the course of antiviral treatment [12–15]. The fact that these mutations are not naturally prevalent in avian strains could suggest that they are not advantageous to the virus, unless a selective drug pressure is applied [34, 40, 41]. There is current widespread resistance of AIV subtypes to the adamantine class of drugs which occlude the M2 ion channel in influenza virus particles inhibiting viral uncoating and thus replication [42]. The establishment of the amantadine resistance in AIV strains is a result of the reported prophylaxis use of these human drugs in birds on poultry farms in South East Asia during the early 2000s when the initial HPAIV H5N1 outbreaks occurred [10]. Unpredictable AIV outbreaks of recent years in poultry which include H7N9, H5N6, H5N2 and H5N8 heighten the risk of a similar practice of poultry prophylaxis with human NAI drugs by poultry owners.

There are 9 different NA subtypes present in influenza virus strains that infect birds (N1-N9) and phylogenetically they fall into two distinct groups: N1-like (N1, N4, N5 and N8) and N2-like (N2, N3, N6, N7 and N9) [43]. The active sites of these two groups of NAs are subtly different as revealed by crystal structure although all four amino acid that were changed in our work were conserved across the two groups and thus our changes (Figure 1) were likely to affect the enzyme activity as they are involved in direct substrate binding (residue 292) and stabilizing framework of the active site (residues 119, 274 and 294) [44, 45]. We observed a significant reduction in relative NA activity for all virus backgrounds when catalytic residue R292 was changed to K (Figure 2). Two framework residue alterations significantly compromised NA activity in five backgrounds: E119V in all except H4N6, and N294S in all except H6N1. H274Y substitution reduced enzyme activity only in two viruses - H5N2 and H7N9, both in the N2-like NA group. Interestingly, we found that the same mutation significantly enhanced NA activity in H5N8 background (Figure 2). Increases as well as decreases in sialidase activity can disrupt the functional HA/NA balance of sialic acid attachment and hydrolysis required by influenza viruses.

We screened in total 30 recombinant viruses, including wt NA, for their resistance to NAIs (Table 2). PR8 was used as a reference strain in each experiment to monitor the inter-assay variability, and its NAI susceptibility profile was consistent with the one reported by others [46]. The resistance pattern for each mutation varied upon strain and antiviral used and thus it is not possible to define functional NAI resistance purely via observation of a specified motif change in the NA sequence. We identified two mutations conferring triple NAI resistance in particular NA subtypes: E119V in H5N8 and H6N1 as well as R292K in H5N6 and H5N2, and the level of resistance varied depending on inhibitor (Table 2; Figure 3 and 4). Such multi-drug resistance have previously been described for E119V in recombinant human H1N1 [33, 47] as well as for R292K in human H3N2 [33] and avian H4N6 subtypes [48]. Our findings have also confirmed multi-drug resistance of R292K variant in H4N6 (ZAN and PER), with predicted high resistance to OSE (precise IC_50_ not determined due to no fluorescence signal change in response to drug). We also found that two NAI resistance-associated substitutions led to the same outcomes across all six virus backgrounds. The E119V mutation conferred moderate resistance to OSE, presumably resulting from the loss of hydrogen bond between the inhibitor and side chain of mutated residue 119 which interacts with oseltamivir in a similar conformation in both N1-like and N2-like groups of NA structure [49]. In contrast all variants carrying H274Y remained sensitive to ZAN which is in line with the published literature suggesting that mutation at position H274, although leading to a conformational change of the active site of the NA, does not disturb the side chain contacts of the C6 of the zanamivir structure unlike oseltamivir and peramivir which have a hydrophobic side chain at this position [18, 32, 37, 50].

R292K substitution affects hydrogen bonding within the active site of NA, leading to reduced interaction with the carboxylate group of NAIs and lowering binding affinity of the drugs [51, 52]. Additionally, binding of OSE to R292K mutant is impeded *via* a second mechanism, where the altered residue restricts conformational changes at position 276 and prevents formation of a hydrophobic pocket required to accommodate the bulky side chain of inhibitor [35, 36, 51, 52]. Along with this structural evidence we demonstrate that R292K signature confers a high resistance to OSE in all virus strains, although the determination of precise IC_50_ values was not possible as the highest drug concentrations used had no impact on fluorescence levels in MUNANA assay (Table 2) or plaque size in MDCK cells (Figure 4 A, D, G and J).

We also compared the drug resistance profiles of avian N1 and N2 tested here with the ones observed in human strains. With one exception (R292K) the resistance patterns of human and avian N1 greatly overlapped - E119V mutation conferred triple drug resistance whereas H274Y and N294S showed reduced inhibition by OSE and PER [16, 17, 20, 33, 37, 47, 53]. To date there have been no reports of R292K naturally occurring in human H1N1 isolates, however a recombinant pH1N1 was moderately resistant to OSE [32]. The same study reported that R292K in avian H5N1 showed no resistance to any of the NAIs [32], whereas our data indicate that R292K in avian H6N1 conferred high resistance to OSE and PER. Similarly to N1, the NAI susceptibility profiles of human and avian N2 were consistent to a major extent, with the exception of N294S which conferred resistance to OSE in human H3N2 [12], whereas the same mutation in avian N2 (represented by H5N2 in this study) led to reduced inhibition by ZAN. We have also shown that the NAI resistance pattern of H7N9 was entirely consistent with the one reported by others [15, 21, 54, 55].

The MUNANA assay routinely used to determine the IC_50_ value of influenza viruses to NAI drugs relies on NA activity knockdown by the drugs, however as we highlight in our study mutation in the NA gene can compromise NA activity in the absence of drug and thus MUNANA assay cannot always be reliably used to determine resistance. We used normalisation of NA activity to haemagglutinin titre (HAU) in this study since this gives us a measure of the amount of HA in a sample. In parallel we performed normalisation to PFU in MDCK cells in this study since reduced NA activity still led to efficient replication in this cell type and thus accurate determination of viable virus in the sample (as determined by plaque assay). (Supplementary Figure 1) which revealed the same trend in NA activity to the HA normalised data set in this case. Changes in NA that alter the sialidase activity often result in concurrent changes in HA or NA that alter the sialic acid binding avidity and thus haemagglutination potential of red blood cells.

We investigated the effect that NAI resistance mediated by the selected mutations had on viral fitness of two subtypes H5N6 and H6N1 in chicken primary kidney cells (cKCs). Despite functionally compromised NAs, the selected NAI-resistant mutants of both H5N6 and H6N1 replicated to comparable or even higher titres than the corresponding wt NAI sensitive virus (Figure 5A and B). Influenza viruses generally produce cell-free virions which then can infect distant cells or tissues in an infected individual, to do this the sialidase function is critical, NAIs prevent this and thus act to inhibit virus dissemination in and from the infected host. Growing in cell culture, viruses may directly spread to the adjacent cells by exploiting intercellular connections reducing the requirement for effective sialidase activity for spread, this has been demonstrated by virus replication even in the presence of NAI drugs [56]. This mechanism may be favoured *in vitro* in case of NAI-resistant mutants with compromised NA activity, thus we did include a NAI sensitive wt virus pre-incubated with OSE in our cKC growth curves and this did knock down virus replication significantly suggesting that compromised NA activity would result in a fitness cost in the context of growth in these cells (Figure 5A & B). However to confirm that the growth patterns we observed in cKC were not a result of NA-independent virus spread, we also performed the growth curves *in ovo* (Figure C-E). We found no significant differences in virus growth between wt H5N6 and NAI-resistant variants (Figure 5 C – E) suggesting no defect in viral fitness in replication *in ovo*. The biological complexity of an embryonated egg as a model for virus fitness assessment presents a suitable link between *in vitro* and *in vivo* work, although assessment in a full chicken model would be required to understand any compromise in virus spread within and between hosts.

It has been shown sequence changes that cause disruption of HA interaction with sialic acid affect NA activity [57] and virus susceptibility to NAIs [58]. On the other hand, NA deficiency can be compensated by readjusting receptor specificities [59, 60] and binding affinities of HA [61, 62]. The HA sequences of our NAI-resistant H5N6 viruses which, despite compromised NA activity, exhibited competitive fitness levels *in vitro* and *in ovo* showed that upon rescue and first passage in eggs, two of the viruses had concurrent HA sequence changes from the wt H5N6 HA (Figure 6A). These two potentially compensatory changes in HA are: Y98F (H3 numbering) in the H5N6 R292K virus reported to diminish receptor binding affinity in human H3N2 viruses [39] and A189T in the H5N6 N294S virus also shown to reduce sialic acid avidity in human H1N1 backgrounds [19]. Indeed we can show that both mutations also significantly reduced the receptor binding avidity of the H5N6 viruses when compared to wt (Figure 6B). The rapid appearance of these HA mutations as consensus sequence changes following rescue and one passage in eggs, of the NA mutant viruses suggests that in the absence of these mutations viable virus carrying the prescribed NA mutations (R292K or N294S) in the H5N6 context would not have been present. In contrast we did not identify any compensatory signatures in HA of H5N6 E119V although its NA activity as well as the receptor binding avidity were significantly reduced. Acquisition of concurrent mutations in HA is the most common mechanism for compensating NA deficiency however this may also be achieved by changes in HA glycosylation pattern occurring near receptor binding site (RBS) [63], altered number of HA units per virion or induced HA steric effects [62]. In several NA subtypes there also exists a second sialic acid binding site, formed of conserved amino acids 367S, 370S, 372S, 400N and 403W, that is distinct from the main sialidase active site and without sialidase function, this site is often termed the NA hemabsorption site [64, 65]. The H5N6 strain we used here is likely to contain such a hemabsorption site and thus modulation of NA structure through drug resistant mutation may alter the NA sialic acid binding potential of the mutant virus and this could be responsible for the reduced RBC avidity of the E119V mutant.

We have shown that the resistance pattern of avian influenza viruses is very diverse and depends on the strain as well as the antiviral used, and there is no universal signature conferring resistance to all NAI drugs in all genetic backgrounds. We have also demonstrated that despite significantly compromised NA activities drug-resistant viruses were able to re-balance the sialidase deficiency by acquiring concurrent changes in HA after just one passage in eggs. Those HA alterations significantly lowered the receptor binding avidities of drug-resistant viruses and restored the fine balance between HA and NA activities. Our study highlights the very difficult-to-foresee molecular pattern of antiviral resistance in influenza viruses and shows how quickly drug-resistant strains can adapt to restore their fitness, similarly to what had been observed worldwide in human H1N1 after 2007 [66]. Our data also showcase the importance of including resistance-associated signatures in the avian surveillance programmes and the need for continuous research aiming to identify novel drug-resistance mechanisms in AIV, as such information would be invaluable for predicting the emergence of future pandemic strains.

## MATERIALS AND METHODS

### Cells, viruses and plasmids

Madin-Darby Canine Kidney cells (MDCK; ATCC CCL-34) and human embryonic kidney cells HEK 293T (ATCC CRL-3216) were maintained in Dulbecco’s modified Eagle’s medium (DMEM) supplemented with 10% fetal bovine serum (FBS; Life Science Production) and 1% penicillin/streptomycin (Gibco). Primary chicken kidney cells (cKC) from specific-pathogen-free (SPF) 2-3 week old Rhode Island Red chicken(s) were prepared in house at The Pirbright Institute (TPI) as described elsewhere [67] and maintained in Eagle’s Minimal Essential Medium (EMEM; Sigma) with Earle’s salt, 2 mM L-glutamine, 2.2 g/L sodium bicarbonate; supplemented with 0.8% bovine serum albumin (BSA; Sigma), 2.95 g/L tryptose phosphate broth (TPB; Sigma) and 1% penicillin/streptomycin. All cell types were incubated at 37°C with 5% CO_2_.

All viruses carrying H5 of HPAIV and/or NA modified to increase the resistance to licensed NAIs were handled in the SAPO4/ACDP3 high containment facility of TPI by trained and authorised personnel.

Neuraminidase (NA) and haemagglutinin (HA) genes of the following strains were synthesised by GeneArt (Invitrogen) and cloned into a pHW2000 vector [68]: H5N6 A/chicken/Jiangxi/02.05 YGYXG023-P/2015 (**<u>GISAID accession nos. EPI661558 and EPI661559</u>**); H5N8 A/scarlet_ibis/Germany/AR44-L01279/2015 (**<u>GISAID accession nos. EPI624533 and EPI624535</u>**); H5N2 A/goose/Taiwan/01031/2015 (**<u>GenBank accession nos. KU646887 and KU646885</u>**); H7N9 A/Anhui/1/2013 (**<u>GISAID accession nos. EPI439509 and EPI439507</u>**); H6N1 A/chicken/Taiwan/67/2013 (**GenBank accession nos. KJ162862 and KJ162860**); H4N6 A/chicken/Hunan/S1267/2010 (**<u>GenBank accession nos. KU160821 and KU160819</u>**).

Single nucleotide changes associated with NAI-resistance were introduced into the NA genes using QuikChange™ site-directed mutagenesis protocol (Stratagene; LaJolla, USA). All recombinant viruses were generated using an eight plasmid reverse genetics system [68] with 6 internal genes (PB2, PB1, PA, NP, M, NS) originated from H1N1 A/Puerto Rico/8/1934 (PR8) and matching HA/NA set from a chosen strain (Table 1). HAs originating from HPAIV retained the multi-basic cleavage site in the HA coding sequences. Viruses were propagated in 10-day old SPF embryonated hen’s eggs (VALO BioMedia GmbH) and allantoic fluid was collected at 48 h (HPAIV) or 72 h (low pathogenic viruses; LPAIV) *post* inoculation. The HA activity of produced virus stocks (haemagglutinating units, HAU/ml) was measured by standard haemagglutination assay [69] with chicken red blood cells (cRBC), and the infectious titres (plaque forming units, PFU/ml) were determined by plaque assay on MDCK cells.

### NA activity and NA inhibition assay

NA activities of recombinant AIV strains were measured using the fluorogenic substrate MUNANA (2’-[4-Methylumbelliferyl]-alpha-D-N-acetylneuraminic acid; Biosynth and Sigma-Aldrich), as described previously [70]. After enzymatic conversion the fluorescent signals corresponding to the amount of released 4-methylumbelliferone (4-MU) were quantified in a microplate reader (Victor X2, Perkin Elmer) with the excitation and emission wavelengths of 355 and 460 nm, respectively. NA activities were expressed as the amount of MUNANA converted to 4-MU within 1 hour at 37°C per 1 HAU of virus stock, and the values were then normalised to the reference strain – PR8, which was run on every plate analysed.

NA inhibition assays were performed using the NA inhibitors: oseltamivir carboxylate (OSE; MedChem Express), zanamivir (ZAN; Sigma-Aldrich) and peramivir (PER; SelleckChem). The final drug concentrations in assay ranged in 4-fold dilutions from 0.015 to 4000 nM. Viruses of each subtype were diluted to reach equivalent levels of NA activity (0.5 – 2 nmol 4-MU/ml*h; varied between strains). Drug concentrations effectively supressing viral NA activity by 50% (IC_50_) were determined using dose-response curve in Graph Pad Prism software (GraphPad Prism version 7.00, La Jolla California USA, www.graphpad.com). Virus drug resistance was classified according to the WHO recommendations (https://www.who.int/influenza/gisrs_laboratory/antiviral_susceptibility/en/) based on the fold increase in IC_50_ compared to the susceptible reference strain (wt NA): normal inhibition (NI) <10-fold; reduced inhibition (RI) 10 to 100-fold; highly reduced inhibition (HRI) >100-fold.

### Virus titration by plaque assay

Infectious virus from harvested allantoic fluid or cell supernatant was titrated by plaque assay on MDCK cells as described before [71], with modified overlay medium: 0.6% agarose (Oxoid) or 0.6 - 0.8% Avicel® Microcrystalline Cellulose and Sodium Carboxymethylcellulose (FMC BioPolymer) in EMEM supplemented with 0.3% BSA, 2.5 mM L-glutamine, 0.2% sodium bicarbonate, 10 mM HEPES, 0.01% DEAE-Dextran, 2 μg/ml TPCK trypsin (all Sigma-Aldrich) and 1% penicillin/streptomycin (Gibco).

### Plaque reduction assay

To determine the effect of NAI drugs on virus spread in cell monolayer we performed plaque reduction assay (PRA) on MDCK cells [72]. Equal volumes of virus inoculum containing between 5 and 100 PFU were added to confluent cells in 12-well plates. After 1 h adsorption at 37°C virus suspension was removed and cells were cultured with 0.6% Avicel overlay supplemented with NAIs: OSE, ZAN or PER. The final drug concentrations in medium ranged in 4-fold dilutions from 0.04 nM to 10 μM; each condition in duplicate. After 72 h incubation at 37°C cells were fixed and stained with 0.1% crystal violet containing 20% methanol. The images of plates were acquired using waterproof digital camera and the diameters of 10 plaques per condition were measured using ImageJ software [73]. The data was analysed in Excel (MS Office) and IC_50_ values were calculated using GraphPad.

### Multi-cycle virus growth curves *in vitro* and *in ovo*

For growth curve *in vitro*, cKC cells in 6-well plates were inoculated with virus in cKC medium at an MOI of 0.001 (4000 PFU/ml), each virus in triplicate. After 1 h at 37°C virus inoculum was removed, cells were washed with PBS and cKC maintenance medium was added to the plates (2 ml/well). A 500 μl aliquot of supernatant was collected from each well at 8, 24, 48 and 72 h *post* infection, and 500 μl of fresh growth medium was added back to the well. To determine the inhibitory effect of NAI on virus growth in cKC, cells infected with wild-type virus were cultured in the presence of 10 µM OSE. All samples were stored at −80°C and titrated by plaque assay on MDCK cells. The growth curves *in vitro* were repeated at least twice.

For growth curve *in ovo* 10-day old embryonated hen’s eggs were inoculated into the allantoic cavity with 100 PFU (1000 PFU/ml) of virus in PBS supplemented with 1% penicillin/streptomycin and 0.35% BSA. Five eggs per each virus and time point were incubated at 37°C and sampled at 10, 24 or 48 h post inoculation by collecting 0.5 - 1 ml aliquot of allantoic fluid. To measure the inhibitory effect of NAI on virus growth *in ovo*, 100 PFU of wt H5N6 virus was pre-incubated with 100 μM PER for 45 min at 37°C, prior to inoculating into eggs. To monitor any adverse effects of drug on embryo survival, three eggs were inoculated with 100 μM PER in virus diluent. All samples were stored at −80°C and titrated by plaque assay on MDCK cells.

### Receptor binding avidity assay

To investigate possible differences in receptor binding avidity of recombinant viruses we performed HA assay with partially de-sialylated cRBCs as described elsewhere [74], with modifications. Briefly, 10% cRBCs in PBS were pre-treated with recombinant neuraminidase from *Clostridium perfringens* (New England Biolabs) for 1.5 hour at 37°C. The final enzyme concentrations ranged in 2-fold dilutions from 1 to 500 U/ml. The erythrocytes were then washed once with PBS and re-suspended to reach a concentration of 1% [v/v]. Virus HA activities were determined on untreated cRBCs following 1 h incubation at 4°C. 4 HAU in 50 μl of each virus was added to 50 μl of sialidase-treated cRBCs solution and incubated for 1 h at 4°C. The highest enzyme concentration allowing for full virus haemagglutination was recorded [U/ml] and expressed as a relative receptor binding avidity.

### Sanger sequencing of viral RNA

Viral RNA (vRNA) was extracted and purified from allantoic fluid using QIAmp Viral RNA kit (Qiagen) according to manufacturer’s instructions, and viral cDNA was synthesised using Verso cDNA reverse transcription kit (ThermoFisher Scientific), influenza universal primers [75] and template RNA. The HA and NA influenza segments (complete or fragments) were amplified from cDNA template using subtype-specific set of primers and purified with QIAquick PCR Purification kit (Qiagen). Nucleotide sequences of viral genes were determined using BrilliantDye™v3.1 Terminal Cycle Sequencing Kit (Nimagen), strain-specific primers and Applied Biosystems 3730 DNA Analyzer.

### Cell viability assay

To reveal any potential cytotoxic effects of antiviral compounds, we seeded MDCK and cKC in 96-well opaque-walled plates and cultured for 72 h in the presence or absence of NAI drugs. Relative cell viability was then quantified using CellTiter-Glo® Luminescent Assay (Promega) according to manufacturer’s protocol.

### Statistical analyses

Statistical analyses were performed in Graph Pad Prism software (GraphPad Prism version 7.00, La Jolla California USA, www.graphpad.com) using one-way ANOVA and Dunnett’s multiple comparisons tests (* P<0.05; ** P<0.01; *** P<0.001; **** P<0.0001).

## FUNDING INFORMATION

This work described herein was funded by BBSRC grant BB/M023362/1. The funders had no role in study design, data collection, data interpretation, or the decision to submit the work for publication.

## ACKNOWLEDGEMENTS

We would like to acknowledge members of the Influenza Virus and Avian Influenza groups at The Pirbright Institute who supported this work; Dr Munir Iqbal, Dr Sushant Bhat, Dr Jean-Remy Sadeyen, Dr Pengxiang Chang and Dr Klaudia Chrzastek.

## AUTHOR CONTRIBUTIONS

The work was conceptualised by HS and DB. Experimental work was executed by DB. The manuscript was written and edited by DB and HS.

---

**Supplementary Figure 1. NA activity profile of recombinant strains normalised to infectivity.** NA activities of six virus subtypes: H5N6 (A), H5N8 (B), H5N2 (C), H7N9 (D), H4N6 (E) and H6N1 (F) were determined using MUNANA assay. Equivalent NA activity levels were chosen for a panel of 5 viruses representing each subtype and normalised to PFU of virus stock. The ratio NA activity/PFU for a reference strain (PR8) was set to 1. The Y axis shows log_10_ fold difference of NA activity/PFU between PR8 and tested viruses. Data represent mean of 3 independent experiments ± SD; statistical significance was determined by one-way ANOVA and Dunnett’s multiple comparisons tests (* P<0.05; ** P<0.01; *** P<0.001).

**Supplementary Figure 2. Relative cell viability of MDCK and cKC cultured in the presence of NAIs.** MDCK (A) or cKC (B) cells were seeded in 96-well black-walled plates and cultured in the presence of NAI drugs for 72 h: (A) 0.15, 0.6, 2.5 or 10 μM of OSE (black bars), ZAN (grey bars) or PER (light grey bars); (B) 0.3, 1.25, 5 or 20 μM OSE, followed by cell viability assessment using luminescent assay (RLU). Untreated cells are shown as white dotted bar, and dashed line on the graph indicates background levels (no cells) also displayed by the white bar with diagonal stripes.

**Supplementary Figure 3. Plaque morphology of wt and NAI-resistant viruses on MDCK cells.** MDCK cells were infected with a panel of H5N6 (A), H5N8 (B), H5N2 (C), H7N9 (D), H4N6 (E) and H6N1 viruses (F) and cultured for 72 h with agarose (A, C and E) or Avicel® overlay medium (B, D and F). Viral plaques were visualised with crystal violet staining.

**Supplementary Table 1.**
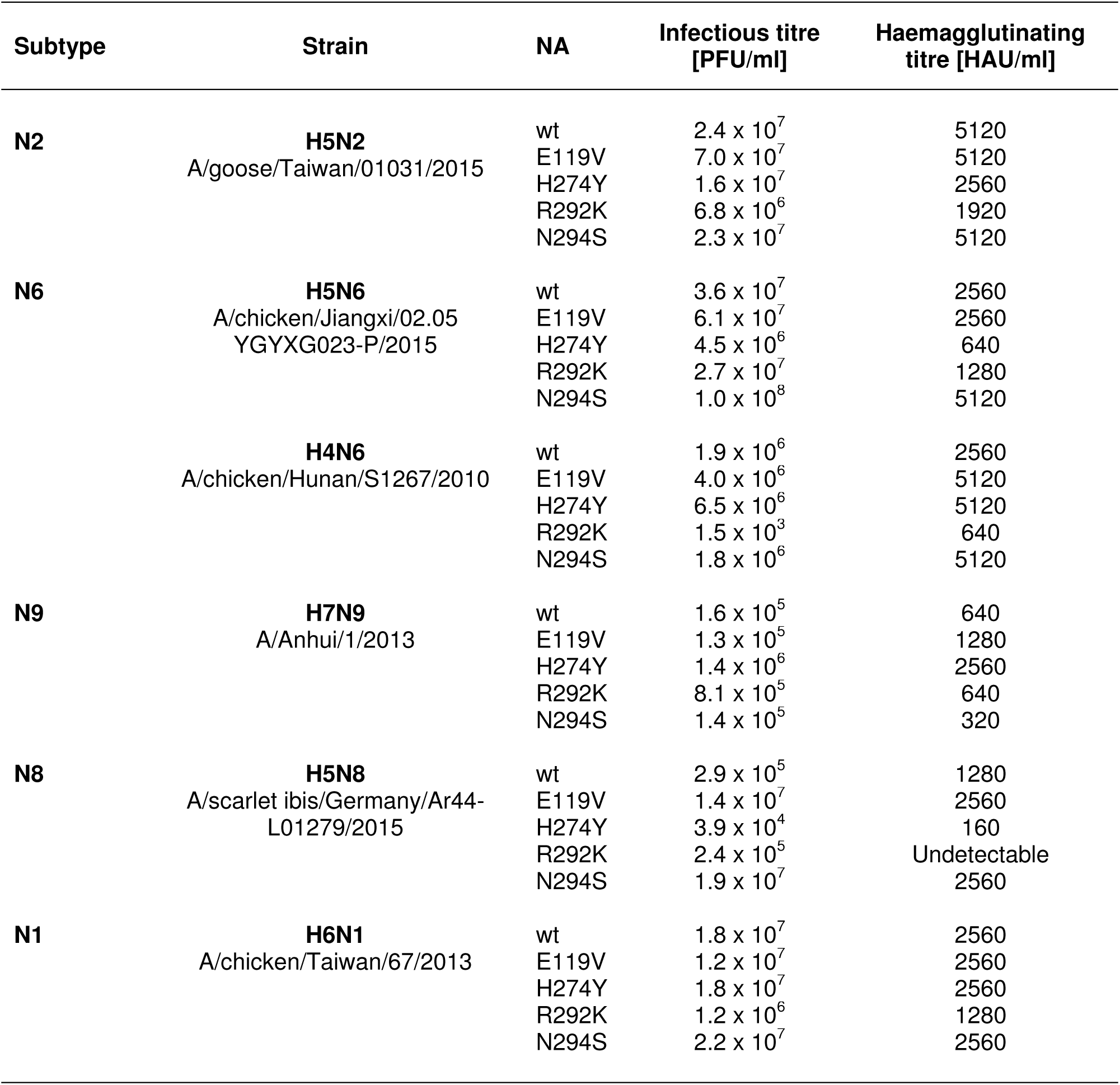
Infectious [PFU/ml] and haemagglutinating [HAU/ml] titres of recombinant viruses carrying wt or mutated NA (P1 egg stock).

